# Agentic cryo-EM structure determination with CryoPilot

**DOI:** 10.64898/2026.10.29.762908

**Authors:** Pranav Punuru, Thomas Klose, Daisuke Kihara

**Affiliations:** Department of Biological Sciences, Purdue University, West Lafayette, IN 47906, USA; Department of Computer Science, Purdue University, West Lafayette, IN 47906, USA

## Abstract

Single-particle cryo-electron microscopy (cryo-EM) relies on mature computational tools, but experts must still inspect intermediate results, choose among alternative processing routes, and decide when earlier choices should be revised. Here we present CryoPilot, a system of two language-model agents that carries cryo-EM data from raw micrographs through three-dimensional reconstruction to validated atomic models. Rather than executing a predefined pipeline, its reconstruction agent evaluates intermediate images and quantitative diagnostics, selects and parameterizes CryoSPARC jobs, tests alternative strategies, compares their outcomes, and retains, revises, or reverts decisions accordingly. A persistent programming environment and session database preserve the complete processing history, making each decision traceable and allowing multi-day workflows to proceed with minimal human intervention. Its modeling agent determines map handedness, builds atomic models with CryoZeta, removes regions the density does not support, and optimizes each model by testing refinement steps against map fit, geometry, and DAQ, a measure of local model–map agreement. Across nine public EMPIAR datasets, eight reconstructions reached 2.61–3.51 Å resolution, and refined models from seven maps agreed with reference structures to within 0.4–4.9 Å Cα RMSD. For a challenging α-latrocrustotoxin dataset, a single change of particle-picking strategy that we suggested enabled CryoPilot to improve the reconstruction from 5.9 to 4.15 Å. CryoPilot establishes an adaptive and auditable framework for agent-driven cryo-EM structure determination.

## Introduction

Single-particle cryo-electron microscopy (cryo-EM) has become a major method for determining three-dimensional (3D) structures of proteins and macromolecular assemblies, with a rapidly growing number of reconstructions deposited in the Electron Microscopy Data Bank (EMDB) ^1^. In single-particle analysis, the microscope records two-dimensional (2D) projection images of many copies of a particle in micrographs. Recovering a three-dimensional (3D) density map from these noisy projections requires a sequence of computational steps, including motion correction, contrast transfer function (CTF) estimation, micrograph curation, particle picking and extraction, 2D classification, initial 3D reconstruction and classification, refinement, and map validation ^2^. Subsequently, atomic model building and model-to-map validation then convert the reconstructed density into an interpretable molecular structure ^3,4^.

Many of the computational steps in the cryo-EM data processing workflow are implemented in mature software such as CryoSPARC ^5^ and RELION ^6^. These programs execute individual processing operations, but researchers must still decide which step to perform next, how to configure its parameters, and whether the resulting output is satisfactory. This decision-making requires expertise because choices made at one stage can strongly affect downstream analysis. For example, particles discarded during 2D classification cannot contribute their views later; a population that appears homogeneous in 2D may be heterogeneous in 3D; and a correction that improves one metric can degrade another. Researchers therefore work iteratively, running one or a few jobs, inspecting micrographs, class averages, map slices, and diagnostic plots, and revisiting earlier decisions when the outcome is unsatisfactory. Although individual computations can run unattended, the overall analysis often pauses for human review. As a result, a single dataset can occupy a specialist for days, and the rationale behind many processing choices is rarely documented beyond the recorded job parameters.

This bottleneck is particularly limiting for less experienced researchers and in high-throughput settings, including structure-based drug discovery ^7^ and cryo-EM facilities that process many datasets. An adaptive system that can evaluate intermediate results, revise its strategy, and document its decisions could reduce dependence on continuous expert supervision, keep multi-step analyses moving without continuous human attention, and allow researchers to oversee multiple datasets in parallel.

Adaptive automation is increasingly needed across science and industry, where complex workflows require not only execution of individual tasks but also evaluation of intermediate results and decisions about what to do next. Advances in artificial intelligence, particularly large language model (LLM) agents, are beginning to enable this capability. Tool-using agents can alternate between action and interpretation ^8^, while code-writing agents can compose computational tools and reuse intermediate results ^9^. Such systems have begun to plan and assess biomedical and chemical research ^10–13^ and to direct automated laboratories ^14^. Cryo-EM data processing is particularly well suited to this form of adaptive automation because its workflow comprises multiple interdependent steps, much of the evidence is visual, decisions must integrate images with quantitative diagnostics, and the consequences of a choice may not become apparent until several processing steps later.

Here we describe CryoPilot, a system of two LLM agents. Its reconstruction agent adaptively processes cryo-EM data from raw movies or micrographs through 3D reconstruction, and its modeling agent then evaluates map contour and handedness, builds macromolecular complexes directly from the reconstructed maps with CryoZeta ^15^, uses DAQ ^16^ to assess residue-level support from the density, and removes unsupported regions before refining the model with Phenix ^17^ and Rosetta ^18^. The researcher provides the input data, acquisition information, and a short scientific goal, but no processing route needs to be specified in advance. The reconstruction agent selects and parameterizes CryoSPARC jobs, inspects intermediate outputs, compares alternative branches, diagnoses unexpected results, and retains, revises, or reverts decisions according to the evidence. It runs one or a few jobs at a time so that each step is informed by preceding results and resumes processing when jobs finish without requiring continuous researcher supervision. The complete session, including submitted jobs, inspected evidence, and agent– researcher exchanges, is stored to make the analysis traceable and recoverable. Because such projects can span dozens of jobs and multiple candidate particle populations, exceeding what a language model can reliably maintain in a single context ^19^, the reconstruction agent therefore adopts a recursive language model (RLM) design ^20^. Project state and processing history are maintained in a persistent Python environment and session database that the model can query with code, while bounded subtasks can be delegated to recursive calls.

We evaluated CryoPilot on nine public EMPIAR datasets spanning a high-symmetry assembly, two enzymes, one bound to a drug-like compound, membrane proteins, an antibody complex, a glycoprotein, a preferred-orientation specimen, and a large flexible toxin, with inputs ranging from movies to pre-extracted particles. Except for two cases, in which we revised the goal statement or particle-picking strategy after an initial run, the input data, acquisition information, and a short goal statement were the only human inputs. Eight reconstructions reached 2.61–3.51 Å resolution; the most difficult dataset, α-latrocrustotoxin, reached 4.15 Å after we supplied a different particle-picking strategy. Across the runs, CryoPilot selected reconstructable classes, designed control refinements, compared competing corrections, detected an input-unit error, and reverted changes that failed to improve the map. For seven reconstructions, the modeling agent, given a goal rather than a procedure, produced refined structures within 0.4–4.9 Å Cα RMSD of reference structures, choosing its own map, a starting structure model, edits, and refinement strategy and optimizing each model: the agent tested refinement programs, restraints, and density weights, kept a change only when map fit, geometry, and residue-level density support improved together, and reverted changes that made the model worse. These results show that language-model agents can assume much of the iterative decision-making that currently keeps experts in the cryo-EM processing and modeling loop while preserving a complete record of how each result was obtained.

## Results

### CryoPilot Overview

CryoPilot consists of two LLM agents (Fig. 1). The reconstruction agent processes cryo-EM data into a 3D reconstruction by executing a sequence of CryoSPARC jobs, evaluating the output of each job, and deciding the next processing step. The modeling agent then builds an atomic model from the reconstructed map and optimizes its agreement with the density. At the start of a project, the researcher provides the input files, acquisition parameters, approximate particle size, expected symmetry, and a short goal statement. The reconstruction agent then selects and configures each CryoSPARC job, waits for it to finish, inspects the result, and decides what to do next; no complete processing route is defined in advance. For focused subtasks, such as reviewing a set of classes, it can also launch a child agent that works in the same project and returns its result to the main agent (Fig. 1a).

**Figure 1:**
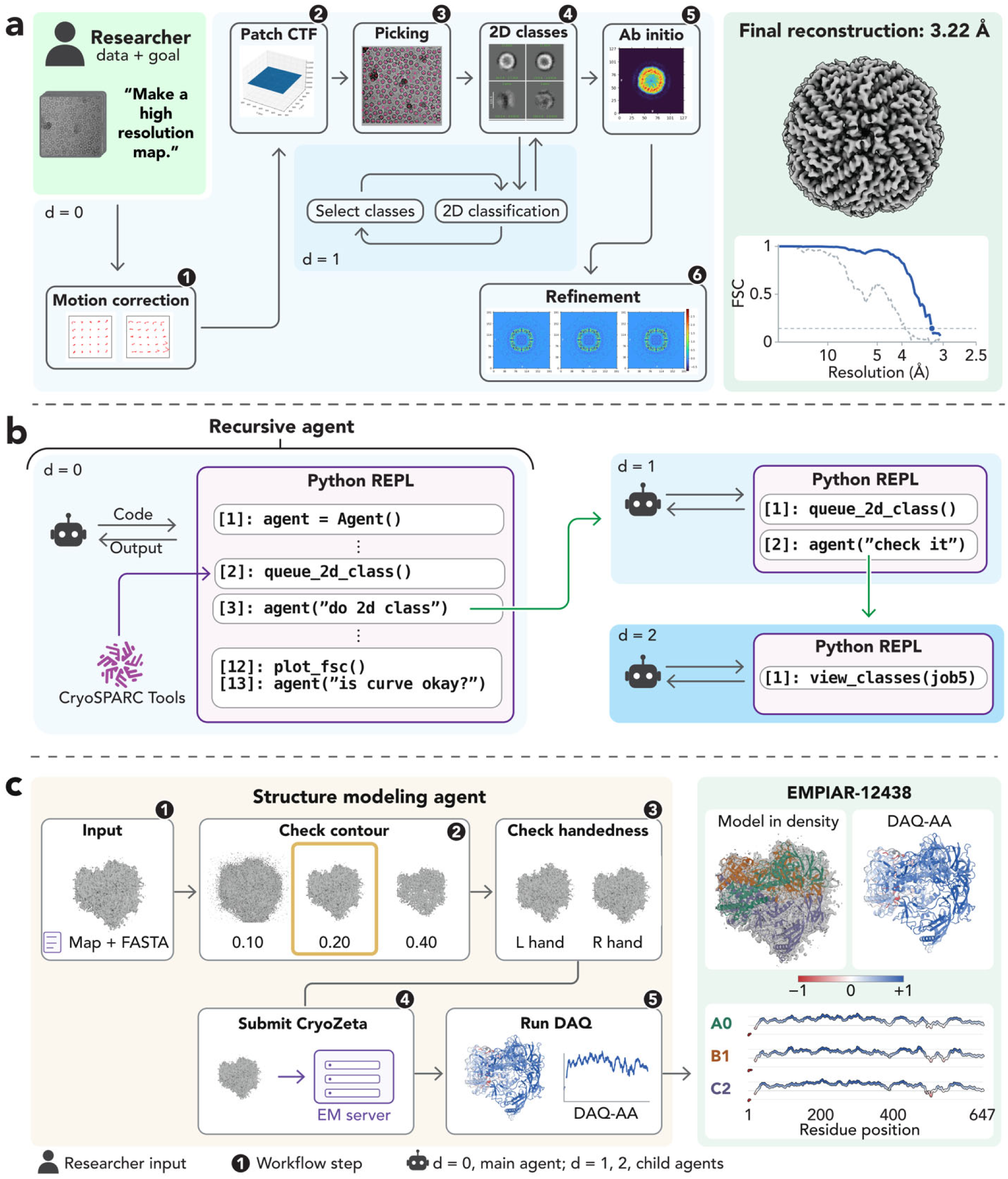
CryoPilot performs recursive agent-based cryo-EM processing and atomic modeling. **a,** Schematic of the cryo-EM reconstruction workflow. Starting from raw movies or micrographs, the reconstruction agent performs motion correction, CTF estimation, particle picking, 2D classification, ab initio reconstruction, and refinement. Rather than following a fixed pipeline, the agent inspects intermediate results and determines the next processing step; the example highlights iterative 2D classification and class selection delegated to a child agent. The workflow is illustrative and does not represent a single experimental run. The person icon marks the researcher input (data and a short goal statement). The FSC curve is the apoferritin validation FSC shown in Fig. 2 (EMPIAR-10146, P40/J29). **b,** The reconstruction agent operates through a persistent Python read–evaluate–print loop (REPL) connected to CryoSPARC. The main agent (depth, d=0) writes and executes code, receives job outputs, and can launch child agents to perform focused subtasks. Child agents can in turn invoke deeper agents (d=1, 2), while operating within the same persistent environment and CryoSPARC workspace. **c,** Atomic-modeling workflow. Given a reconstructed map and protein sequences, the modeling agent evaluates contour level and map handedness, submits the selected map to CryoZeta for atomic model building, and evaluates residue-level model–map agreement with DAQ. The example shown is EMPIAR-12438 (P31/J29). The final model is displayed within the cryo-EM density and colored by chain, together with DAQ-AA coloring and chain-specific DAQ-AA profiles; the red–white–blue scale ranges from −1 to +1. d = 0 marks the main agent and d = 1, 2 its child agents, and numbered circles give the order of the workflow steps in a and c.

The reconstruction agent is built using a recursive language model (RLM) design ^20^. The language model acts by writing Python code, which is executed in a persistent Python environment throughout the project (Fig. 1b; Algorithm 1, Methods). This environment provides access to the CryoSPARC workspace, a cryo-EM processing guide compiled from the CryoSPARC tutorials and case studies ^21^, the full conversation history, and the function used to launch child agents. Because the environment persists between steps, information such as job identifiers, particle counts, and intermediate analyses remains available as the workflow progresses. Images such as class averages, map slices, and diagnostic plots are shown to the model only when it explicitly loads them for inspection.

As a project grows, active conversation may become too long for the model to use efficiently. The reconstruction agent therefore keeps the most recent interactions in full and summarizes older ones (Algorithm 2, Methods), while preserving the complete record in the Python environment and an SQLite database. This allows the agent to recover information even after context compression or an interrupted session. The same records also make the workflow auditable: for each decision, we can trace the images and numerical results the agent examined, the action it took, and the outcome of the resulting job.

The modeling agent builds atomic models from reconstructed maps (Fig. 1c). It receives the map, its resolution, and the sequences and copy numbers of the assembly, and works through a set of tools rather than a fixed pipeline. It first assesses the map, choosing the map version and a contour level from the enclosed volume and the connectivity of the density, and tests the map hand by fitting an experimental structure of the same or a homologous protein into the map and its mirror image with VESPER and comparing the fit scores of the two hands. It then uses CryoZeta ^15^ through its API, which returns five candidate models from the map, and scores each candidate by map‒model correlation, geometry, and DAQ, which estimates residue-level agreement between the model and density ^16^. It also checks the assembly against the construct for misconnected chains and for segments that are absent from the mature protein, such as signal peptides and cleavage linkers. We use CryoZeta because it builds models of protein and nucleic-acid complexes directly from cryo-EM maps, performs strongly in benchmark evaluations ^15^, and is accessible through a server API. DAQ, Phenix ^17^, and Rosetta ^18^ run locally on the compute node hosting the modeling agent. The agent then removes residues that the density does not support and optimizes the model by real-space refinement in Phenix and density-guided relaxation in Rosetta. For each round it chooses the program to use, restraints, and density weight, keeps the round only when map fit, geometry, and DAQ improve together, and otherwise returns to the previous model. Rather than following a fixed sequence of steps, it is guided by instructions that specify the modeling goal and the types of evidence an experienced modeler would consider. The final output is a refined atomic model annotated with regions of strong and weak density support, together with a decision log that records each choice, its evidence, and the alternatives considered.

Both agents use the general-purpose language model GPT-5.5 through its API. We developed the software harness that controls what information is provided to the model and enables it to interact with CryoSPARC. We also designed the agent instructions and tools and, for the reconstruction agent, a searchable cryo-EM processing guide, refining these components through repeated test runs. The reconstruction agent was developed using one CryoSPARC tutorial dataset, EMPIAR-10025, and the modeling agent in earlier runs on maps from this study. The agents themselves run on CPUs; GPUs are used only by CryoSPARC and the structure-modeling programs (Methods).

### Summary of 3D map reconstruction

We applied CryoPilot to nine datasets from EMPIAR, the public archive of raw cryo-EM images ^22^, selected to cover common processing challenges, such as junk-rich micrographs, conformational heterogeneity, preferred orientation, and flexibility (Extended Data Table 1) ^23–31^. Each CryoPilot run began with a goal statement that generally asked the agent to produce the best-supported reconstruction while evaluating alternative processing strategies, such as different particle-picking methods, 2D class selections, or refinement approaches. The datasets comprised a high-symmetry tutorial dataset (apoferritin); a tetrameric enzyme, pyruvate kinase M2 (PKM2), bound to the drug-like Compound 5; a tetrameric membrane channel (TRPA1); an asymmetric antibody complex (HER2–Fab); a heavily glycosylated trimer (HIV-1 Env); a second tetrameric channel, TRPV1, in lipid nanodiscs with bound toxin; a tetrameric enzyme with preferred orientation (aldolase); a specimen with severe preferred orientation (hemagglutinin); and a large, flexible, asymmetric toxin (α-latrocrustotoxin). The assemblies range in molecular mass from 140 to 689 kDa. The projects started from movies, micrographs, or extracted particles, so the runs entered the processing pipeline at different points (Fig. 2). Eight datasets have deposited reconstructions in the EMDB (EMD-10575, 6267, 18188, 51753, 8117, 76791, 8731, and 13642), with reported resolutions ranging from 2.95 to 4.24 Å; the apoferritin dataset has no corresponding EMDB entry and is a tutorial dataset used in cisTEM ^23^.

**Figure 2:**
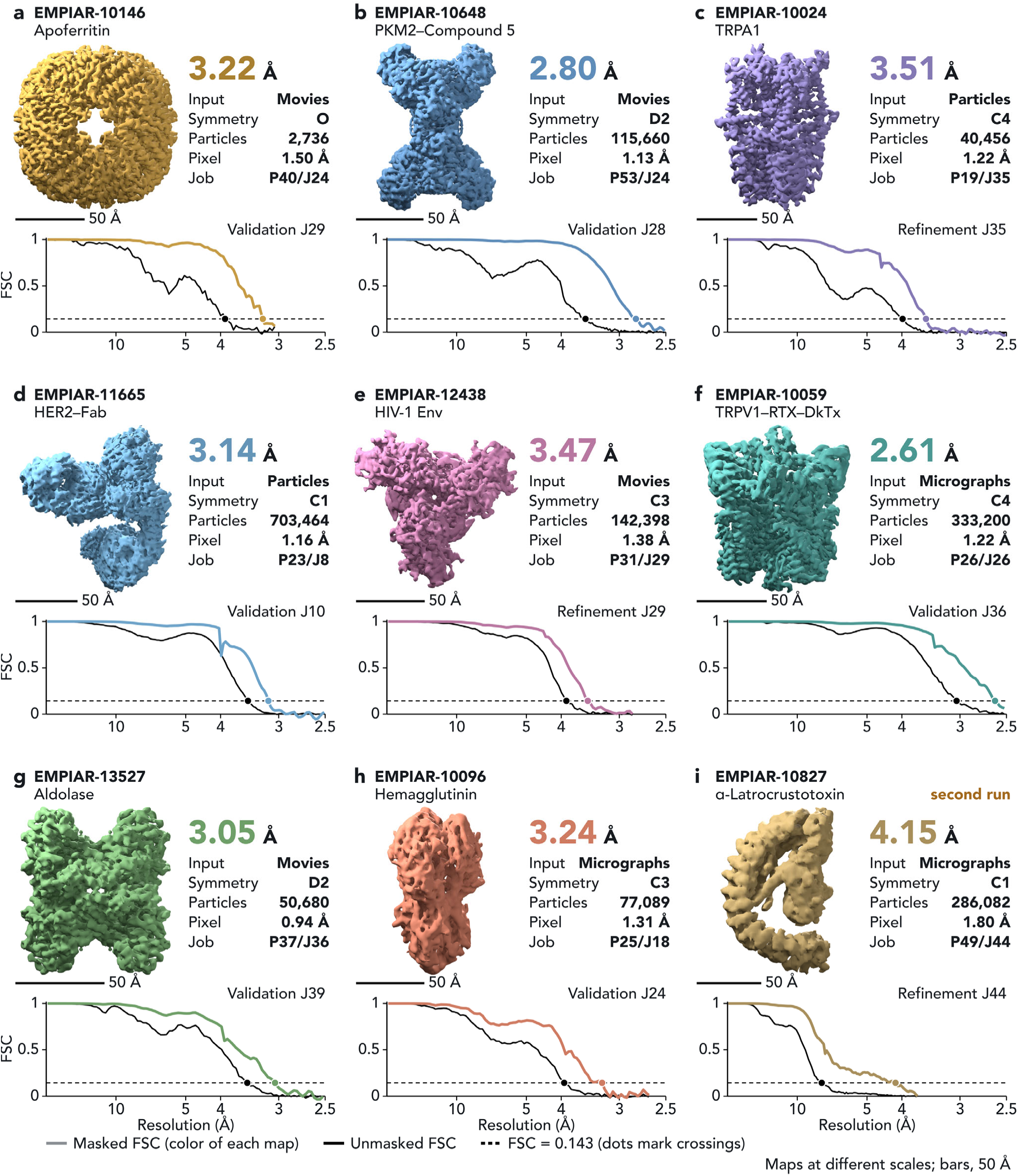
CryoPilot reconstructions of nine EMPIAR datasets. For each dataset, CryoPilot received the deposited movies, micrographs, or particles, the acquisition parameters, and a short goal statement, and processed the data in CryoSPARC without a predefined workflow. Each panel shows a map reconstructed by CryoPilot, labeled with its EMPIAR accession and specimen: **a**, apoferritin; **b**, PKM2–Compound 5; **c**, TRPA1; **d**, HER2– Fab; **e**, HIV-1 Env; **f**, TRPV1–RTX–DkTx; **g**, aldolase; **h**, hemagglutinin; **i**, α-latrocrustotoxin. Each panel gives the map, its global resolution, the type of starting data, the imposed symmetry, the number of particles accepted by the refinement, the pixel size, and the job, above the gold-standard FSC redrawn from the CryoSPARC output, which has two curves. The masked FSC (color of the map; noise-substitution corrected where CryoSPARC reports it) defines the reported resolution. The unmasked FSC (black) is calculated without a mask and is therefore lower. The dashed line marks FSC = 0.143, and dots mark where the two curves cross it. Where the agent ran a final Validation (FSC) job to check the resolution of its chosen map, the panel shows that curve, which reuses the refinement half maps and mask; the other panels show the refinement curve after mask auto-tightening. Maps are ordered as in Extended Data Table 1 and shown at different scales; bars, 50 Å. The EMPIAR-10827 map shown is from the second run (P49/J44), which followed a particle-picking strategy that we suggested and started from exposures corrected in the first run.

Eight of the nine reconstructions reached global resolutions of 2.61–3.51 Å at FSC = 0.143 ^32,33^, whereas the ninth, α-latrocrustotoxin, reached 4.15 Å after two runs (Fig. 2). For the second run, we instructed CryoPilot to refine its particle picking on micrographs screened with Micrograph Junk Detector ^34^ and denoised with Micrograph Denoiser ^35^. For HIV Env, we also revised the goal statement after a first run (below). These resolutions are comparable to those reported in the original studies (Extended Data Table 1). For six datasets, our numerical resolution estimates are better than the deposited values, but we do not interpret this as evidence of superior maps because CryoSPARC has improved since several of the original studies, masking and FSC procedures differ across studies, and the deposited hemagglutinin map was reconstructed from tilted data. Rather, the comparison shows that CryoPilot reaches the resolution range achieved by expert processing of the same datasets. Resolution also did not simply track particle number: TRPV1 reached 2.61 Å from 333,200 particles, the octahedral apoferritin reached 3.22 Å from only 2,736 particles, and PKM2 reached 2.80 Å from 115,660 particles, fewer than the 287,873 particles used for the deposited reconstruction.

How CryoPilot proceeded depended on the dataset. For apoferritin, PKM2, TRPA1, and HER2– Fab (Fig. 2a–d), it followed largely conventional workflows of 15–37 jobs. For datasets starting from movies, it performed motion correction, CTF estimation, blob picking, and 2D classification, followed by ab initio reconstruction and homogeneous and non-uniform refinement. For apoferritin, it additionally re-picked particles using templates generated from the initial 2D classes. For datasets starting from imported particles, it reconstructed and refined the deposited particle sets; for TRPA1, it further removed heterogeneous particles using 2D and 3D classification and heterogeneous refinement. Even in these cases, the agent made dataset-specific choices of the type normally made by an expert. For PKM2, for example, it retained a broader 2D selection than two stricter alternatives to preserve rare views, enlarged the extraction box to account for signal delocalization, and performed ab initio reconstruction and heterogeneous refinement without symmetry before imposing D2 symmetry on the dominant class. It then applied global and local CTF refinement and reference-based motion correction ^36^, re-refining after each step and improving the resolution from 3.52 to 3.39, 2.93, and finally 2.79 Å.

For the five more difficult datasets, HIV Env, TRPV1, aldolase, hemagglutinin, and α-latrocrustotoxin (Fig. 2e‒i), CryoPilot went beyond a standard workflow by forming hypotheses about what was limiting the reconstruction and designing additional jobs to test them, resulting in runs of 29– 55 jobs. For HIV Env (Fig. 2e), an earlier run using a general goal statement and no specified symmetry produced a strongly anisotropic map at 6.9 Å resolution. We therefore revised the goal statement to highlight the substantial junk in the micrographs, specify C3 symmetry, and ask CryoPilot to compare alternative particle-picking strategies while carrying multiple particle sets forward. In the revised run, CryoPilot added a second picking branch using denoised micrographs, retained both strict and broad 2D selections, and then selected among four classes generated by heterogeneous refinement. For TRPV1 (Fig. 2f), it tested whether a minority cluster identified by 3D variability analysis represented a distinct structural state by refining that cluster separately as a control. For aldolase (Fig. 2g) and hemagglutinin (Fig. 2h), which showed strongly non-uniform orientation distributions, it quantified the viewing directions and tested particle rebalancing strategies. For α-latrocrustotoxin (Fig. 2i), the 2D selection of the first run could not separate genuine views of this flexible, preferentially oriented toxin from junk, and the map stopped at 5.98 Å. We therefore suggested a revised particle-picking strategy, which CryoPilot tested on a pilot subset before applying it to the full dataset.

CryoPilot treated each correction as a hypothesis to be tested rather than a step to be accepted automatically. It applied per-particle (local) CTF refinement ^37,38^ in nine runs and retained it only when the subsequent refinement showed a clear benefit. In seven runs, apoferritin, TRPA1, two TRPV1 runs, hemagglutinin, aldolase, and the first α-latrocrustotoxin run, the agent reverted to the uncorrected particles. In the five cases examined in detail, the re-refined map was unchanged or worse; where the session record included this check, CryoPilot had first verified that the per-particle defocus changes were centered near zero (Fig. 3b, 3c). By contrast, for PKM2, local CTF refinement improved the resolution from 3.39 to 2.93 Å and was retained, while for HER2–Fab the agent kept a refinement that combined global and local corrections. It treated global aberration corrections similarly, retaining them only when the following refinement improved the reconstruction.

**Figure 3:**
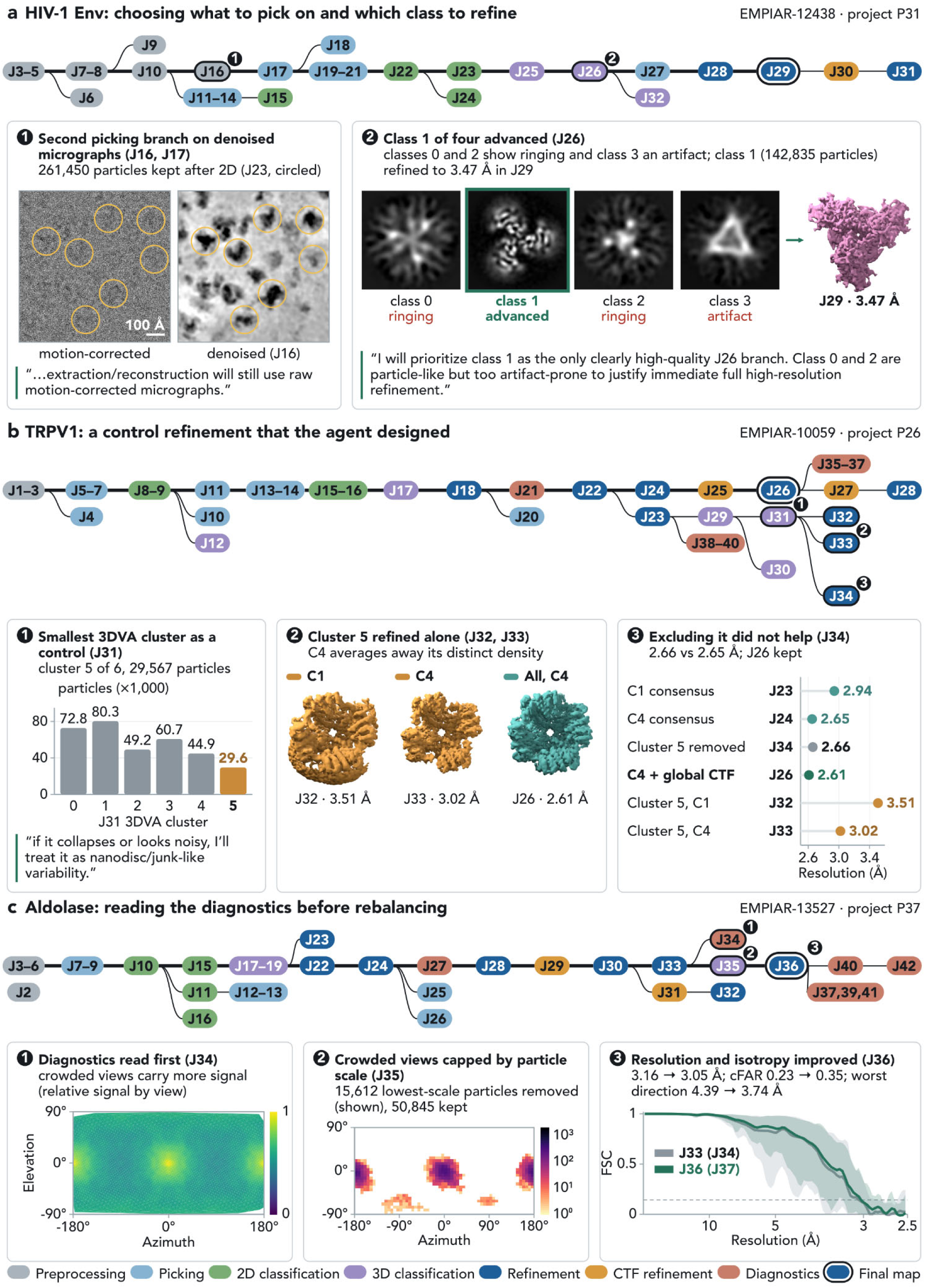
Examples of decisions made by CryoPilot from intermediate results. For three representative projects, the CryoSPARC job tree is shown above panels that illustrate selected decision points. Each node represents a job, is labeled with its CryoSPARC job number, and is colored by job type. Each job is positioned to the right of its most recent input job, so branches indicate alternative processing paths explored by the agent. The thick line traces the lineage leading to the final map, whose job node is ringed. Consecutive unbranched jobs of the same type are merged into a single node labeled with the job-number range; failed jobs are omitted. Numbered nodes indicate the decision points described below, with the corresponding numbered panels showing the evidence considered by the agent. Quotations are the agent’s own statements from the session records. a, HIV-1 Env (EMPIAR-12438, P31; 30 jobs). (1) J16: the agent denoised the micrographs and picked a second particle set from them. The same micrograph region is shown before and after denoising, with particles retained after 2D selection (J23) circled. (2) J26: central slices of the four classes from heterogeneous refinement, together with the agent’s assessments. Class 1 was advanced and refined to 3.47 Å (J29). b, TRPV1 (EMPIAR-10059, P26; 39 jobs). (1) J31: particle counts for the six 3DVA clusters; the agent selected the smallest cluster as a control. (2) J32 and J33: the selected cluster refined without and with C4 symmetry, shown as top views at a common scale alongside the all-particle consensus map (J26). (3) J34: global resolutions of the compared maps. Removing the cluster did not improve the consensus reconstruction. c, Aldolase (EMPIAR-13527, P37; 36 jobs). (1) J34: relative signal by viewing direction from orientation diagnostics (heat map derived from the CryoSPARC plot, with axes redrawn). (2) J35: viewing directions of the 15,612 particles removed during rebalancing, which capped each of 192 bins at 427 particles by removing the lowest-scale particles first. (3) J36: directional FSC before (J33; diagnostics J34) and after (J36; diagnostics J37) rebalancing. Lines show the mean across 3,072 cones, and shaded bands show the 1st–99th percentiles.

The following sections focus on the five difficult datasets, highlighting representative decision points where CryoPilot interpreted intermediate results, changed its processing strategy, and tested whether those changes improved the reconstruction.

### HIV Env: selection of a reconstructable 3D class and detection of a parameter-unit error

Fig. 3 shows the job trees of three projects in which the agent made notable decisions. Each tree shows how the agent branched, compared alternatives, and abandoned branches that did not help, and the steps highlighted in the text are shown with their evidence below each tree.

The first example is EMPIAR-12438 (HIV-1 Env), for which CryoPilot obtained a 3.47 Å map in 30 jobs (Fig. 2e). After motion correction, CTF estimation, junk detection, and exposure curation (J3– J10), the agent performed blob picking on the motion-corrected micrographs, followed by 2D classification (J11–J15). Because this branch contained many small junk particles, CryoPilot opened a second branch: it denoised the micrographs (J16), performed blob picking with a larger particle-size setting (J17–J22), and extracted the particles from the original micrographs (Fig. 3a, step 1). It reviewed all 200 2D classes in ten batches and retained both strict and broad selections (J23 and J24; 261,450 and 407,072 particles) rather than committing to one. It then performed ab initio reconstruction (J25) and heterogeneous refinement (J26) on the strict set.

Heterogeneous refinement of the strict set (J26) produced four markedly different volumes (Fig. 3a, step 2). Classes 0, 2, and 3 showed compact density surrounded by concentric ringing, whereas class 1 showed discrete, protein-like density. CryoPilot therefore advanced class 1 (142,835 particles), which refined to 3.47 Å (J27–J29). It subsequently rejected global CTF refinement because the resolution changed by less than 0.002 Å and the conical FSC area ratio (cFAR), a measure of directional isotropy, decreased from 0.84 to 0.80; the agent therefore retained the uncorrected J29 map (J30–J31; Extended Data Fig. 1a).

The same run also illustrates error detection. While attempting to reject exposures with more than 35% junk area, CryoPilot supplied 0.35 to a parameter expressed in percent ^39^. The job kept only the 1,568 of 8,482 exposures with at most 0.35% junk area, without reporting an error. From this implausible result, the agent recognized the parameter mismatch, resubmitted the job with a threshold of 35, and retained 7,961 exposures (J9–J10; Extended Data Fig. 1b). This shows how inspecting job outputs allowed the agent to identify and correct an input error.

### TRPV1: a control refinement of a minority cluster and continuous peripheral variability

The second example is EMPIAR-10059, TRPV1 in lipid nanodiscs bound to resiniferatoxin and double-knot toxin (DkTx) ^26^, for which CryoPilot obtained a 2.61 Å map in 39 jobs (Fig. 2f). After particle picking, 2D classification, ab initio reconstruction, and refinement (J1–J22), the agent generated C1 and C4 consensus maps (J23–J24). It retained global CTF refinement, which produced the final C4 map (J25– J26), but reverted local CTF refinement because it provided no further improvement (J27–J28). The final reconstruction closely matched the deposited map, EMD-8117: using the same soft spherical mask for both half-map pairs, our map crossed FSC = 0.143 at 2.91 Å compared with 3.18 Å for the deposition (Extended Data Fig. 2a,b).

CryoPilot then designed a control to test whether a minority population represented a distinct state (Fig. 3b). Among six clusters from 3D variability analysis (J29–J31), it selected the smallest cluster (29,567 particles; step 1), reasoning that coherent refinement would support a genuine minority state whereas incoherent density would suggest junk. The cluster refined to 3.51 Å without symmetry and 3.02 Å with C4 symmetry (J32–J33; step 2). C4 averaging sharpened the channel core but removed the density distinguishing the cluster, indicating that its variation lay mainly in the peripheral DkTx, linkers, and nanodisc. Removing the cluster did not improve the consensus (J34; 2.66 Å versus 2.65 Å; step 3), so CryoPilot retained the all-particle C4 map together with a C1 map (2.94 Å) to preserve asymmetric peripheral density.

In an independent run on the same dataset, 3D variability analysis ^40^ and clustering led the agent to conclude that the heterogeneity was largely continuous and peripheral rather than composed of discrete states. Along the leading variability component, peripheral density rotated around a stable tetrameric core, resembling the relative rotation of cytoplasmic domains reported among TRPV channel structures ^41^.

### Aldolase: signal-aware orientation rebalancing

The third example (Fig. 3c) is EMPIAR-13527, rabbit aldolase, for which CryoPilot obtained a 3.05 Å map in 36 jobs (Fig. 2g). Because the goal statement requested comparison of rebalanced and unrebalanced refinements where justified, CryoPilot first assessed whether rebalancing was warranted. After particle picking, 2D classification, and refinement (J2–J28), it retained global CTF refinement but reverted local CTF refinement (J29–J32), as for TRPV1. Non-uniform refinement then reached 3.16 Å from 66,457 particles (J33). Orientation diagnostics (J34) revealed strong anisotropy, with cFAR = 0.23 and directional resolutions ranging from 2.85 to 4.39 Å ^42^. The agent then examined signal, rather than particle counts alone, across viewing directions. Because the over-represented views also carried stronger signal, it reasoned that random removal could lose more signal than it gained in isotropy (Fig. 3c, step 1). In J35, it therefore capped the most populated viewing-direction bins while preferentially removing particles with the lowest per-particle scale factors ^43^ (Fig. 3c, step 2).

The rebalanced subset retained 50,845 of 66,457 particles and refined to 3.05 Å (J36; Fig. 3c, step 3). Resolution and isotropy improved together: cFAR increased from 0.23 to 0.35, the sampling compensation factor SCF* from 0.87 to 0.96, and the worst directional resolution improved from 4.39 to 3.74 Å ^44–46^ (Fig. 3c, step 3; Extended Data Fig. 1c). By contrast, the more aggressive random rebalancing tested for hemagglutinin (below) reduced resolution. Although the specimens and rebalancing settings differed, these cases suggest that which particles are removed can be as important as whether rebalancing is performed at all.

### Hemagglutinin: retention of rare views under preferred orientation

The next example shown in Fig. 4, hemagglutinin (EMPIAR-10096), illustrates a limitation imposed by the data rather than by a single processing decision. Using only untilted images, CryoPilot obtained a 3.24 Å map in 29 jobs (Fig. 2h). The job tree (Fig. 4a) proceeds from curation and blob picking through a single 2D classification and refinement to the final map, J18, from which the alternative refinements branch out. Using the same half-map FSC procedure for both reconstructions, J18 crossed FSC = 0.143 at 3.77 Å, compared with 5.09 Å for the deposited map EMD-8731 reconstructed from 40°-tilted images (Fig. 4b). The key challenge was to preserve the rare side and oblique views throughout processing.

**Figure 4:**
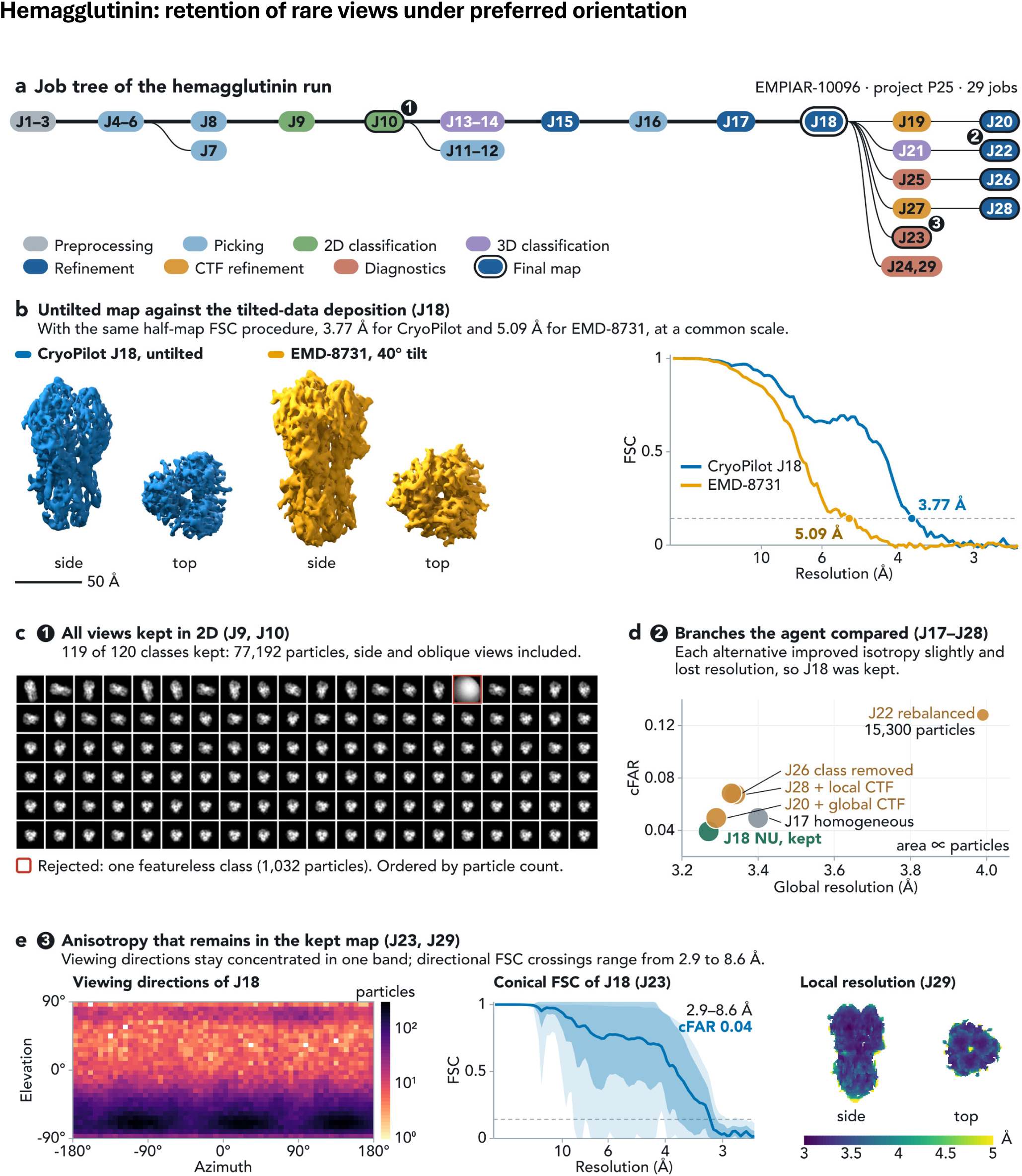
Reconstruction of hemagglutinin under strong preferred orientation from untilted images. Hemagglutinin (EMPIAR-10096, project P25). **a,** CryoSPARC job tree for the 29-job run, drawn as in Fig. 3. Numbered markers indicate the decision points illustrated in **c–e**, and the additional outlined jobs denote alternative branches compared in **d**. **b,** CryoPilot reconstruction (J18, blue) and the tilted-data deposition EMD-8731 (gold; reflected and rigidly aligned for display), shown in matched side and top views at the same scale. The corresponding half-map FSC curves were calculated using the same soft spherical mask procedure; the FSC = 0.143 crossings were 3.77 Å for J18 and 5.09 Å for EMD-8731. **c,** All 120 class averages from 2D classification J9, ordered by particle count in the final iteration from left to right and top to bottom (2,402 to 255 particles per class). CryoPilot rejected only one featureless class (outlined; 1,032 particles) and retained the remaining 119 classes, comprising 77,192 particles (J10), thereby preserving side and oblique views. **d,** Alternative refinement branches tested by the agent, plotted by global resolution and conical FSC area ratio (cFAR); symbol area is proportional to particle count. J18 gave the best global resolution, whereas each alternative modestly improved isotropy at the cost of resolution, so J18 was retained. **e,** Residual anisotropy in the retained J18 reconstruction. Left, viewing-direction distribution of the J18 particles on a logarithmic particle-count scale. Middle, conical FSC analysis from orientation diagnostics J23, showing the mean over 3,072 cones, ±1 s.d. and 1st–99th percentile bands, the range of directional FSC = 0.143 crossings (2.9–8.6 Å), and the cFAR reported by CryoSPARC. Right, local resolution at FSC = 0.5 ^47^ from J29, averaged through the map density in side and top views. Scale bar, 50 Å.

Hemagglutinin strongly prefers top views on the grid, and the original study collected data at a 40° stage tilt to recover missing orientations ^29^. An untilted reconstruction therefore depends on retaining the few side and oblique views, which can resemble junk because they form weak class averages. The goal statement explicitly asked CryoPilot to preserve such views. It curated 447 micrographs to 379, used circular and elliptical blob picking over 75–135 Å to capture elongated side views, and inspected all 120 2D classes. It rejected only one featureless class and retained the other 119, totaling 77,192 particles (Fig. 4c), consistent with the inclusive selection recommended for strongly oriented specimens ^48^.

CryoPilot then improved the map without reducing this population. Using all 77,192 particles, it obtained a C3 consensus at 4.09 Å, re-extracted particles at the original 1.31 Å pixel size when the downsampled map approached its sampling limit, and improved the map to 3.40 Å by homogeneous refinement and 3.27 Å by non-uniform refinement ^44^ (Fig. 4d).

The agent next tested four common strategies for further improvement (Fig. 4d). Global and local CTF refinement produced 3.29 and 3.34 Å maps, respectively, and removing a small heterogeneous class gave 3.33 Å, all worse than the 3.27 Å J18 map. Orientation rebalancing increased cFAR ^42^ from 0.039 to 0.128 but reduced the dataset to ∼15,300 particles and the resolution to 3.99 Å. Because every alternative slightly improved isotropy at the expense of resolution, CryoPilot retained J18 and reported its anisotropy explicitly. The deposited EMD-8731 was reported at 4.2 Å ^29^; because the box sizes differ, the common half-map comparison in Fig. 4b indicates only that the untilted reconstruction is at least comparable; it does not give a precise ranking.

Anisotropy nevertheless remained. Directional FSC crossings ranged from 2.90 to 8.62 Å, cFAR was 0.04, and viewing directions remained concentrated within a narrow elevation band, although local resolution was 3.1–3.8 Å across most of the map density (10th–90th percentile; Fig. 4e). Processing cannot recover views that were never recorded, but CryoPilot preserved the rare views that were present. The modeling agent later determined that this reconstruction had inverted handedness and built its model into the map with the corrected hand (Fig. 6f).

### *α*-Latrocrustotoxin: an unsuccessful first run and a second run with our guidance

The final example, α-latrocrustotoxin (EMPIAR-10827), was a difficult dataset and the only one for which CryoPilot’s initial strategy did not produce a map that the reconstruction agent judged suitable for atomic modeling. The protein is large, curved, flexible, asymmetric, and strongly affected by preferred orientation. The deposited reconstruction, EMD-13642, reached 4.03 Å from 376,474 particles ^30^. CryoPilot’s first run stopped at 5.98 Å, whereas a second run using a particle-picking strategy suggested by us reached 4.15 Å (Fig. 2i; Fig. 5).

**Figure 5:**
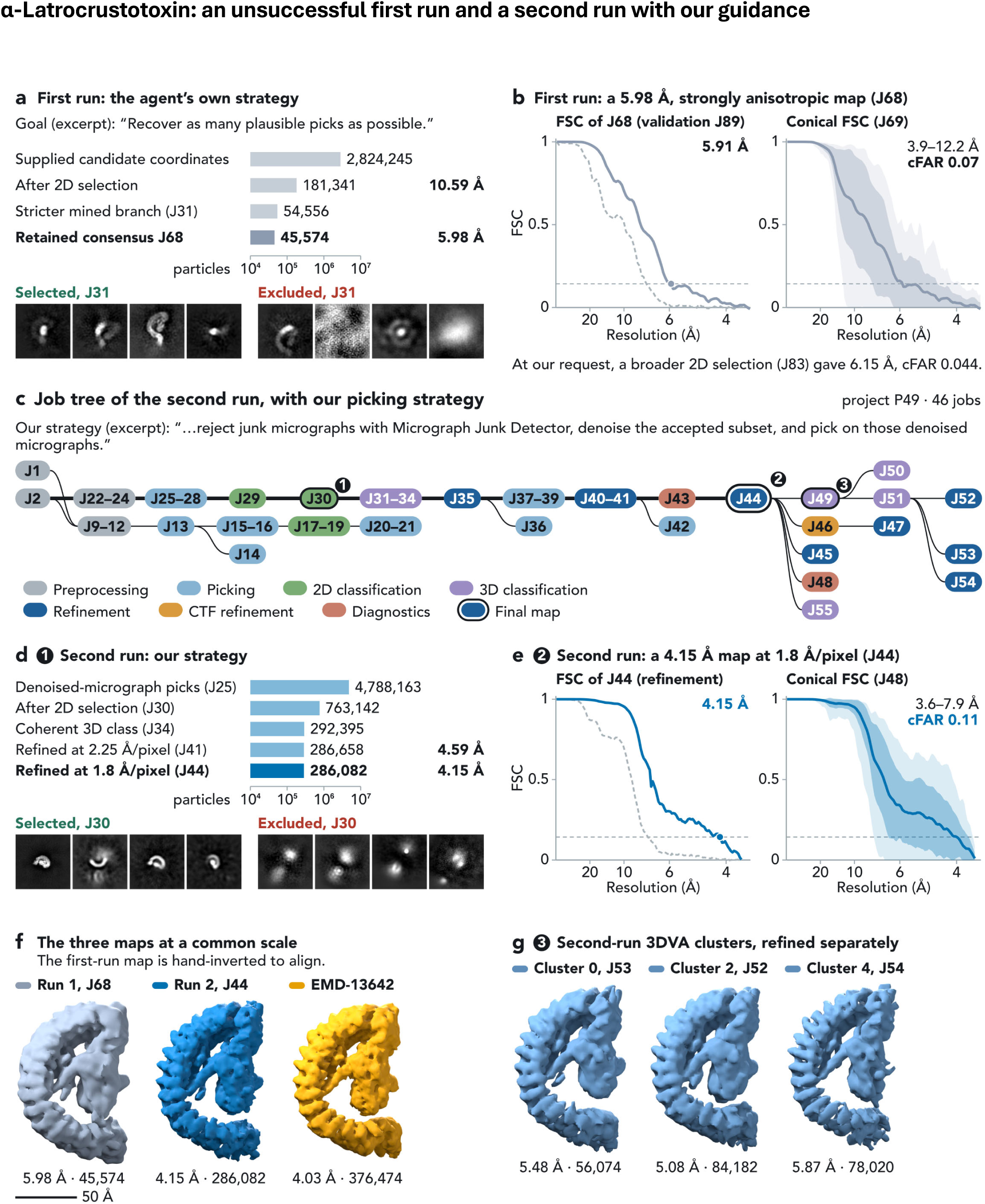
Reconstruction of a highly heterogeneous specimen using two particle-picking strategies. α-Latrocrustotoxin (EMPIAR-10827). **a,** First run (project P39), showing an excerpt from the goal statement, particle counts through CryoPilot’s initial processing strategy on a logarithmic scale, and the first four selected and excluded class averages from 2D selection J31. **b,** First-run consensus map J68. Left, validation FSC from a separate masked calculation (J89; 5.91 Å). Right, conical FSC analysis from J69, showing strong directional anisotropy. Below, the result of a broader 2D selection that we requested (J83). **c,** CryoSPARC job tree for the second run (project P49; 46 jobs), drawn as in Fig. 3. This run started from the corrected micrographs of the first run and followed the revised particle-picking strategy specified by us, excerpted above the tree. Numbered markers indicate the decision points illustrated in **d, e,** and **g**. **d,** Particle counts through the second-run strategy and the first four selected and excluded class averages from 2D selection J30. **e,** Refinement FSC of the second-run consensus J44 (4.15 Å) and its conical FSC analysis from J48, displayed as in Figs. 2 and 4e. **f,** First-run map J68, second-run map J44, and deposited map EMD-13642, aligned to J44 and shown at the same scale. The first-run map could be aligned only after inversion of its hand. **g,** Separate refinements of three well-populated 3DVA clusters from the second run, shown at the same scale. Their similar overall structures, together with additional 3D classification (J55), led the agent to interpret the remaining heterogeneity as predominantly continuous flexibility rather than discrete compositional states. Scale bar, 50 Å.

In the first run, CryoPilot was instructed to recover as many plausible particles as possible, starting from 2,824,245 supplied candidate coordinates and adding independent picking (Fig. 5a). The main difficulty arose during 2D selection: compact, curved particle-like averages appeared among both selected and rejected classes. One branch retained 181,341 of 334,264 particles but reached only 10.59 Å, and later branches reduced the supplied candidates to 54,556 and 66,900 particles before 3D curation. The final consensus, J68, reached 5.98 Å from 45,574 particles (5.91 Å by an independent masked FSC calculation), with cFAR = 0.07 and directional resolution of 3.9–12.2 Å (Fig. 5b).

We then asked CryoPilot to test a broader particle population. A more permissive 2D selection retained 201,687 particles, but subsequent 3D curation produced maps at 6.64 Å from 111,363 particles and 6.15 Å from 67,161 particles (J83), neither better than J68. The larger branch was also more anisotropic (cFAR 0.044 versus 0.067), so the agent retained J68 rather than assuming that more particles would produce a better map (Fig. 5b). It also recognized that its 3D variability clusters were correlated with the gold-standard half-set assignment and therefore treated the cluster maps as exploratory rather than comparing their FSC values. The agent concluded that the ∼6 Å anisotropic map was unsuitable for atomic modeling and did not propose a model. We therefore classified this run as unsuccessful, with the limiting step being recovery of an informative particle population.

For the second run, we specified a different particle-finding strategy: remove heavily contaminated micrographs using Micrograph Junk Detector ^34^, denoise the accepted micrographs ^35,49^, pick on the denoised images, and first test the procedure on a small subset from both collection days. CryoPilot then made the downstream processing decisions. On a 120-micrograph pilot subset (J9–J21), it compared blob and template picking and generated templates from 26 of its own 2D classes. On the full dataset, it performed junk detection, denoising, and template picking (J22–J28), retaining 8,708 of 9,161 exposures, picking 4.8 million candidates, filtering them to 1.8 million, and extracting 1,355,455 particles from the original micrographs (Fig. 5c, d). After reviewing all 200 2D classes (J30) and the rejected classes separately, it restored five rejected classes as plausible rare views and retained 763,142 particles. Heterogeneous refinement against five ab initio references (J31–J34) then isolated a coherent class of 292,395 particles.

At 2.25 Å/pixel, refinement reached 4.59 Å, close to the 4.5 Å Nyquist limit. CryoPilot therefore re-extracted the particles at 1.8 Å/pixel and refined 286,082 particles to 4.15 Å (Fig. 2i; Fig. 5e). For comparison, the first run reached 5.98 Å from 45,574 particles and EMD-13642 reached 4.03 Å from 376,474 particles (Fig. 5f). The first-run map aligned with the other two only after inversion of its hand. Although a subsequent non-uniform refinement gave a nominal 3.67 Å, near the new 3.6 Å Nyquist limit, CryoPilot judged this estimate optimistic because the map remained anisotropic (cFAR 0.11; directional resolution 3.6–7.9 Å) and retained J44 as the consensus map (Fig. 5e).

Finally, CryoPilot examined the remaining heterogeneity using three analyses (Fig. 5g). The 3DVA latent space formed a continuous distribution; three separately refined clusters (5.08–5.87 Å) differed mainly by slight bending; and 3D classification without pose updates produced four classes with the same overall topology. The agent therefore concluded that α-latrocrustotoxin undergoes predominantly continuous flexibility rather than forming discrete structural states. Together, the two runs localize the first failure to particle recovery: the downstream reasoning was appropriate in both runs, but improving the particle population required one strategic intervention from us.

### Atomic models built from the maps and validated against density

CryoPilot’s modeling agent converts each reconstructed map into a refined and validated atomic model. Given the map, its resolution, and the assembly sequences, it is asked to produce the most accurate model supported by the density without following a predefined workflow. In each run, the agent selected the map and contour, determined map hand, chose among CryoZeta candidates, checked the construct, removed unsupported residues, and iteratively refined the model. Refinement steps were evaluated by map fit, geometry, and DAQ, and changes that worsened the model were reverted. Its tools include the CryoZeta server API ^15^, DAQ, map–model fitting, and Phenix and Rosetta (Methods).

We applied the agent to seven assemblies: PKM2, HER2–Fab, HIV Env, TRPV1, aldolase, hemagglutinin, and α-latrocrustotoxin. We also modeled the poorer 5.98 Å α-latrocrustotoxin map from the unsuccessful first reconstruction run, giving eight modeling runs in total. All proceeded without human intervention and completed in 2.6–10.8 h (Fig. 6; Extended Data Tables 2 and 3). Each required 55–79 tool calls and one or two CryoZeta jobs.

**Figure 6:**
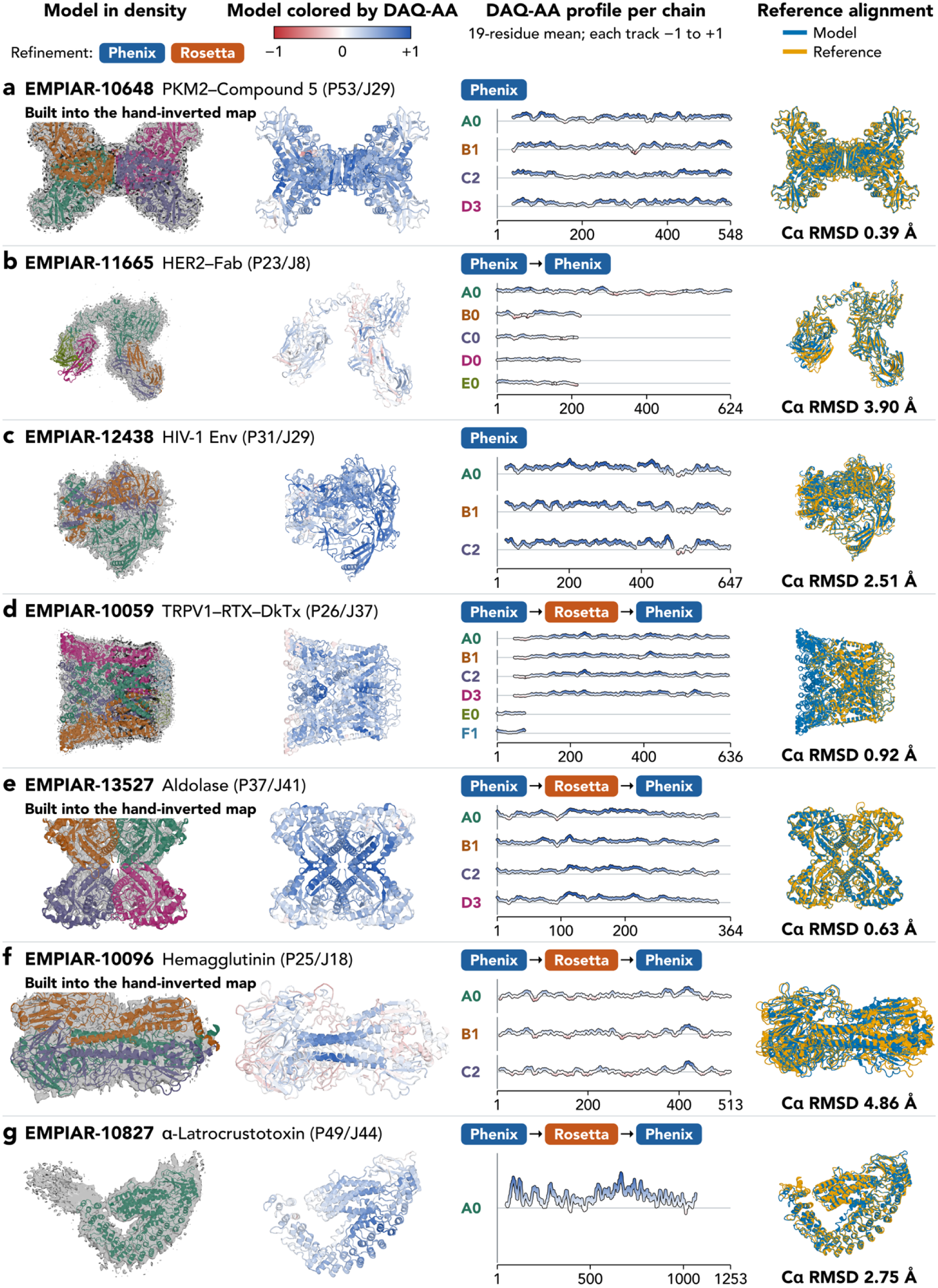
Atomic models built by CryoPilot’s modeling agent, validated against density and reference structures. **a**, PKM2–Compound 5 (EMPIAR-10648; reference PDB 6TTF); **b**, HER2–Fab (EMPIAR-11665; 8Q6J); **c**, HIV-1 Env (EMPIAR-12438; 6UDA, a related structure); **d**, TRPV1–RTX–DkTx (EMPIAR-10059; 5IRX); **e**, aldolase (EMPIAR-13527; 1ADO, a related structure); **f**, hemagglutinin (EMPIAR-10096; 3WHE); **g**, α-latrocrustotoxin, second run (EMPIAR-10827; 7PTX). Each row names the map job and shows, from left to right, the final model in the map density, colored by chain; the same model colored by DAQ-AA; the DAQ-AA profile of each chain; and the whole-complex superposition of the model (blue) on the reference structure (gold), with the Cα RMSD over aligned residues (Extended Data Table 2). The boxes above each profile are the refinement steps that produced the final model, in the order run (Phenix, blue; Rosetta, orange). Rejected steps and the other decisions of the agent are listed in Extended Data Table 3. DAQ-AA estimates how well the density supports each residue: blue (+1), well supported; white (0), no clear support; red (−1), not supported. Colors and profiles use centered 19-residue means, and DAQ-AA was computed at the contour chosen by the agent. Red segments, most visible for HER2–Fab and for the anisotropic hemagglutinin map, are weakly supported regions that the agent kept because the density there is connected (Results). Rows a, e, and f are marked “Built into the hand-inverted map”: the agent modeled these maps in the inverted hand, along z. For HIV Env, gp120 and gp41 were compared as separate chains (Extended Data Table 2). Map isosurfaces are shown at display contours chosen by the authors.

The resulting models agreed closely with experimental reference structures. Whole-complex superpositions gave Cα RMSDs of 0.39 Å for PKM2, 0.63 Å for aldolase, 0.92 Å for TRPV1, 2.51 Å for HIV Env, 2.75 Å for α-latrocrustotoxin, 3.90 Å for HER2–Fab, and 4.86 Å for hemagglutinin over 974– 2,044 aligned residues, with TM-scores ^50^ of 0.87–0.99 (Fig. 6; Extended Data Table 2). Alignments covered 94.5–99.8% of each reference. Because no models were deposited with the Env and aldolase maps, we used closely related structures, PDB 6UDA for Env (99.6% and 100% sequence identity for gp120 and gp41) and PDB 1ADO for aldolase (99.7%), as references.

The agents did not see these evaluation comparisons. Experimental structures could be used only as supporting evidence, for example to determine map hand. In every run, the agent compared fits to the map and its mirror image using VESPER; the chosen hand had a DOT score 1.6–6.1 times higher than the alternative (Extended Data Table 3). Model extent and refinement were instead determined from map– model agreement, DAQ ^16^, and geometric validation. Final models had MolProbity scores of 0.82–1.46 and clashscores of 1.1–8.4 ^51^, which indicate good geometry.

For PKM2 (Fig. 6a), fitting the deposited tetramer PDB 6TTF strongly favored the hand-inverted map (VESPER ^52^ DOT score 5,450 versus 894), so the agent modeled the inverted hand ^32^. CryoZeta built the model into the unsharpened map at a contour matching the expected tetramer volume, and refinement used the sharpened map. The agent removed residues 1–37 of each chain, including the histidine tag, thrombin site, and poorly supported native residues, reducing the clashscore from 50 to 7 and the MolProbity score from 2.65 to 1.39. One Phenix refinement raised CC_mask from 0.77 to 0.84 and improved MolProbity to 1.11. The agent then stopped rather than risk over-regularization with Rosetta. The final model had a Cα RMSD of 0.39 Å.

For HER2–Fab (Fig. 6b), the agent chose a contour that matched the expected complex volume and retained the original hand based on VESPER fitting. Among five CryoZeta candidates, it prioritized higher DAQ and fewer residues outside the density over a negligible difference in CC_mask. It also retained the weak but continuous density of the Fab constant domains. A Phenix refinement introduced nine Cβ deviations, so the agent restarted with stronger geometric restraints, which reduced them to one. Whole-model minimization removed the remaining deviation but reduced CC_mask from 0.57 to 0.48; the agent rejected that result and minimized only the affected residue. The final model had a Cα RMSD of 3.90 Å.

For HIV-1 Env (Fig. 6c), the agent lowered the contour from 0.100 to 0.075 to account for glycan and peripheral density. VESPER strongly favored the original hand (DOT score 1,998 versus 752), so the agent ran CryoZeta only on the original map. It removed the signal peptide, a disordered loop, and the cleavage linker, reducing residues outside the density from 144 to 28 and the clashscore from 19.4 to 8.8. Restrained Phenix refinement increased CC_mask from 0.70 to 0.83. One error remained, which we found during evaluation: CryoZeta connected each gp120 to the gp41 subunit of a neighboring protomer so that the engineered SOS disulfide Cys482–Cys588 linked different chains. The agent removed the unsupported linker but did not reassign the gp41 chains, so we evaluated gp120 and gp41 separately against the reference (Extended Data Table 2). The final Cα RMSD was 2.51 Å.

For TRPV1 (Fig. 6d), the agent chose a contour that included protein together with lipid and nanodisc density. It compared sharpened and unsharpened maps to judge the weak N terminus, removing residues 1–47 but retaining residues 48–87 because connected density was visible in the unsharpened map. It also retained residues 624–629 and removed 630–636. The model also included most of the N-terminal ankyrin-repeat domain, which is absent from the experimental reference but supported by density (mean DAQ-AA 0.28–0.30 per chain, compared with 0.39–0.43 for the rest of the model). After Phenix refinement, the agent rejected a second restrained run because it worsened geometry as judged by MolProbity. Rosetta improved MolProbity to 0.82 but lowered CC_mask to 0.64. B-factor-only refinement then restored CC_mask to 0.81 without changing the coordinates. The final Cα RMSD was 0.92 Å over the aligned reference residues.

For aldolase (Fig. 6e), VESPER fitting of an identical-sequence aldolase structure favored the hand-inverted map (1,309 versus 476), so the agent modeled the inverted hand. Because the five CryoZeta candidates had nearly identical CC_mask values, it selected the model with the best DAQ and MolProbity score. It removed residues 345–364 from each chain because DAQ and overlays showed that they lacked density support. A Phenix run slightly improved CC_mask but worsened MolProbity and clashes, so it was rejected. Rosetta improved MolProbity to 0.82 but reduced CC_mask to 0.71, and a short Phenix refinement restored CC_mask to 0.77 while retaining good geometry. The final model had a Cα RMSD of 0.63 Å.

For hemagglutinin (Fig. 6f), the agent again selected the hand-inverted map based on VESPER fitting, consistent with the deposited EMD-8731 map, which aligns with ours only after reflection (Fig. 4b). This was the most anisotropic reconstruction and produced the largest Cα RMSD, 4.86 Å, and the lowest mean DAQ. The agent retained residues 326–333, a weakly supported segment. It deferred this decision until after refinement and kept the segment because no continuous stretch of residues remained outside the density. It also identified an option that had increased rotamer outliers to 5.97%, reran Phenix without it and with Ramachandran restraints, and reduced the outliers to 1.44%. Rosetta then improved MolProbity from 2.26 to 1.16 and clashscore from 23.1 to 3.7, although CC_mask fell from 0.54 to 0.50. A subsequent Phenix coordinate refinement restored map fit but degraded geometry, so the agent rejected it and refined only B-factors and occupancies.

For α-latrocrustotoxin (Fig. 6g), the agent submitted CryoZeta jobs on both sharpened and unsharpened versions of the 4.15 Å map and selected a candidate from the sharpened-map run based on DAQ. It retained several density-supported segments absent from PDB 7PTX and refined the model with Phenix, Rosetta, and a final B-factor-only Phenix step. As for TRPV1, this restored map fit after Rosetta without moving the coordinates. The final model had a Cα RMSD of 2.75 Å.

One decision in this run was influenced by the reference structure, which the agent had fitted for the hand test. Although density analysis flagged only residues 1188–1253 as unsupported, the agent also removed residues 1067–1187, ending the model at residue 1066, the last residue resolved in 7PTX, despite positive mean DAQ for the removed segment. This truncation was not supported by the density and reduced model coverage. In contrast, when modeling the poorer 5.98 Å map from the first reconstruction run, the agent retained this region as lower confidence and produced a model with a Cα RMSD of 2.99 Å and a TM-score of 0.923 against 7PTX. Its much lower mean DAQ score, 0.069 versus 0.343, reflected the poorer map quality (Extended Data Tables 2 and 3).

## Discussion

We developed CryoPilot, a system of two language-model agents that connects cryo-EM data processing, 3D reconstruction, and atomic modeling in an adaptive workflow. Its main advance is not automation of individual CryoSPARC jobs, but automation of the decisions that connect them. CryoPilot inspects intermediate results, compares alternatives, tests hypotheses, and retains or reverts changes according to the evidence. Across the evaluated datasets, it selected reconstructable particle populations, tested alternative processing strategies, responded to preferred orientation and heterogeneity, detected an input error, and rejected corrections that did not improve the reconstruction. The modeling agent similarly chose map hand and contour, edited models according to density support, and balanced map fit against stereochemical quality rather than following a fixed refinement workflow.

This automation offers several practical advantages. It can keep multi-step analyses progressing without continuous expert supervision, allowing researchers to oversee multiple datasets in parallel. Because every inspected result, decision, and submitted job is recorded, the workflow is auditable and reproducible by other researchers. This may also help standardize processing across users and facilities. For less experienced researchers, the recorded decision process can provide guidance on how expert-style choices are made. These properties may be particularly useful in high-throughput settings such as cryo-EM facilities and structure-based drug discovery.

Two features of CryoPilot are important for this behavior. First, the multimodal model can combine visual evidence, such as micrographs, class averages, map slices, and diagnostic plots, with quantitative results. Second, the persistent programming environment gives the agent access to job parameters, outputs, calculations, and complete processing history. This allows it to compare results quantitatively and maintain coherent projects extending over dozens of jobs without relying entirely on the language model’s active context.

Coupling reconstruction directly to atomic modeling extends the workflow from map generation to an interpretable structural result. Given a goal rather than a prescribed procedure, the modeling agent produced structures within 0.4–4.9 Å Cα RMSD of reference structures and adapted its decisions to each map. The results also show the value of retaining expert oversight. For α-latrocrustotoxin, CryoPilot’s initial particle-selection strategy did not recover a sufficiently informative population, but after we specified a different particle-finding strategy, the agent tested it and completed the remaining processing independently. A practical near-term role for such agents may therefore be that the researcher defines the objective and provides high-level guidance for difficult cases, while the agent performs the detailed execution, testing, and documentation.

One limitation is that the agents did not catch every modeling error automatically. This can be partly addressed by giving the agent more explicit instructions and adding targeted checks for known failure modes, such as chain connectivity, construct consistency, and disulfide pairing. More generally, the agent instructions and guidance can be refined over time. CryoPilot currently has access to a cryo-EM processing guide compiled from CryoSPARC documentation, tutorials, and case studies; a natural next step is to enrich this resource with lessons from previous runs. Such experience-based guidance could help the agent recognize recurring problems and apply successful strategies from past analyses to new datasets.

Cryo-EM software has advanced rapidly at the level of individual processing algorithms, but the decisions connecting those algorithms remain largely manual. CryoPilot shows that a language-model agent with access to visual evidence, computational tools, persistent project state, and its own processing history can make many of these decisions and can incorporate expert direction when needed. This provides a framework for adaptive, auditable cryo-EM structure determination in which the boundary between autonomous processing and expert intervention can be progressively evaluated and refined.

## Methods

### Design for the map reconstruction agent

CryoPilot consists of two language-model agents, a reconstruction agent and a modeling agent. The reconstruction agent processes single-particle cryo-EM data in CryoSPARC. It is implemented as a recursive language model (RLM) harness, a software layer that allows the language model to execute Python code, control CryoSPARC, retain a record of previous steps, and delegate focused subtasks to child agents. The researcher defines the scientific goal, CryoSPARC performs the image-processing calculations, and the language model selects each processing step and revises the workflow as new results become available.

A language model is used through its application programming interface, without additional training or fine-tuning. Its behavior as an agent is determined by the instructions, available tools, and a cryo-EM processing guide. We developed these instructions using a subset of the CryoSPARC tutorial dataset, EMPIAR-10025, which contains the *Thermoplasma acidophilum* 20S proteasome ^53^, and refined them iteratively after each test run. EMPIAR-10025 was therefore excluded from the evaluation datasets. In this study, the agent ran on a CPU node, and CryoSPARC jobs ran on a lane of four workstations with eight GPUs (four NVIDIA RTX PRO 6000 Blackwell and four RTX A6000).

The reconstruction agent has five components (Fig. 1b). We provide brief explanation for each.

#### Python REPL

The language model acts by writing Python code, which is executed in a persistent read– evaluate–print loop (REPL), following the RLM design ^20^. Using code allows the agent to perform small calculations needed for processing decisions, such as counting particles in each class or comparing FSC curves. Because the REPL persists throughout the project, information such as job identifiers, particle counts, and intermediate results remains available for subsequent steps.

#### CryoSPARC client

The REPL contains a CryoSPARC client library^54^ connected to one project workspace. Through it, the agent reaches job types, parameters, job submission and status, outputs, logs, and images. All image processing therefore uses the algorithms of CryoSPARC, and a researcher can inspect and continue any run in the CryoSPARC interface. The agent can list and search all 126 job types in the job register of CryoSPARC v5.0.2 (93 stable and 10 beta). Jobs in the reported projects use 45 of these types.

#### Tools

The language model has three tools. *run_code* runs a code block in the REPL, and *run_and_wait* runs a code block and then waits for the queued CryoSPARC jobs to finish. *view_image* loads an image, such as a sheet of 2D class averages or a diagnostic plot, into the context of the model, because much of the evidence in cryo-EM processing is visual. The instructions tell the agent to run one or a few jobs at a time, so that each decision follows an inspection of the previous results.

#### Processing guide and session database

The REPL also contains a searchable cryo-EM processing guide with practical information on CryoSPARC jobs and parameters, which supplies domain knowledge in place of fine-tuning. An SQLite database records every message and tool result, so that each session can be audited and continued after an interruption. The guide is a markdown file of about 37 kB in ten sections, compiled by language-model summarization of the tutorials and case studies of the CryoSPARC Guide (https://guide.cryosparc.com/processing-data/tutorials-and-case-studies)^21^, and we inspected each summary. The guide covers general decisions, such as when to re-pick particles, how to diagnose preferred orientation, and when to refine CTF parameters, rather than the workflow of any one tutorial. Individual names of biomolecules or database entry identities are not included. When a session starts, the harness loads the guide into the REPL, and the instructions tell the agent to search it and read its sections when it chooses workflows, parameters, diagnostics, or corrective actions.

#### Language model

The reconstruction agent uses GPT-5.5 at the highest (xhigh) reasoning setting ^55^. We use this setting because extended reasoning gives the best performance across the published benchmarks of the model ^56^. We selected this model on the basis of a preliminary 2D class-selection benchmark, in which it scored 90.3–93.2% of the maximum across reasoning settings, compared with 79.0–86.1% for Claude Opus 4.8 (Extended Data Fig. 3). Similar models also drive other recent scientific agents, including the Virtual Lab, which uses GPT-4o ^10^, and a GPT-5-driven autonomous laboratory ^14^. We counted the tokens of each message and image in the input of every model call with the o200k_base tokenizer. A reconstruction run used 13–93 million input tokens and 0.05–0.40 million output tokens, about US$14–106 at list prices (median US$32). These runs took 2–148 h from the first to the last agent step (median 29 h). For the first α-latrocrustotoxin run, we counted only the work up to J90, excluding the idle time before our follow-up request.

#### Agent loop

In each iteration, the language model reads the active context and writes a response. The harness runs each code block in the response and adds only the printed output to the context. When a response contains no code, the loop ends and the text of the response is the answer. Algorithm 1 shows this loop:

##### Algorithm 1 Recursive language-model (RLM) agent with a persistent REPL

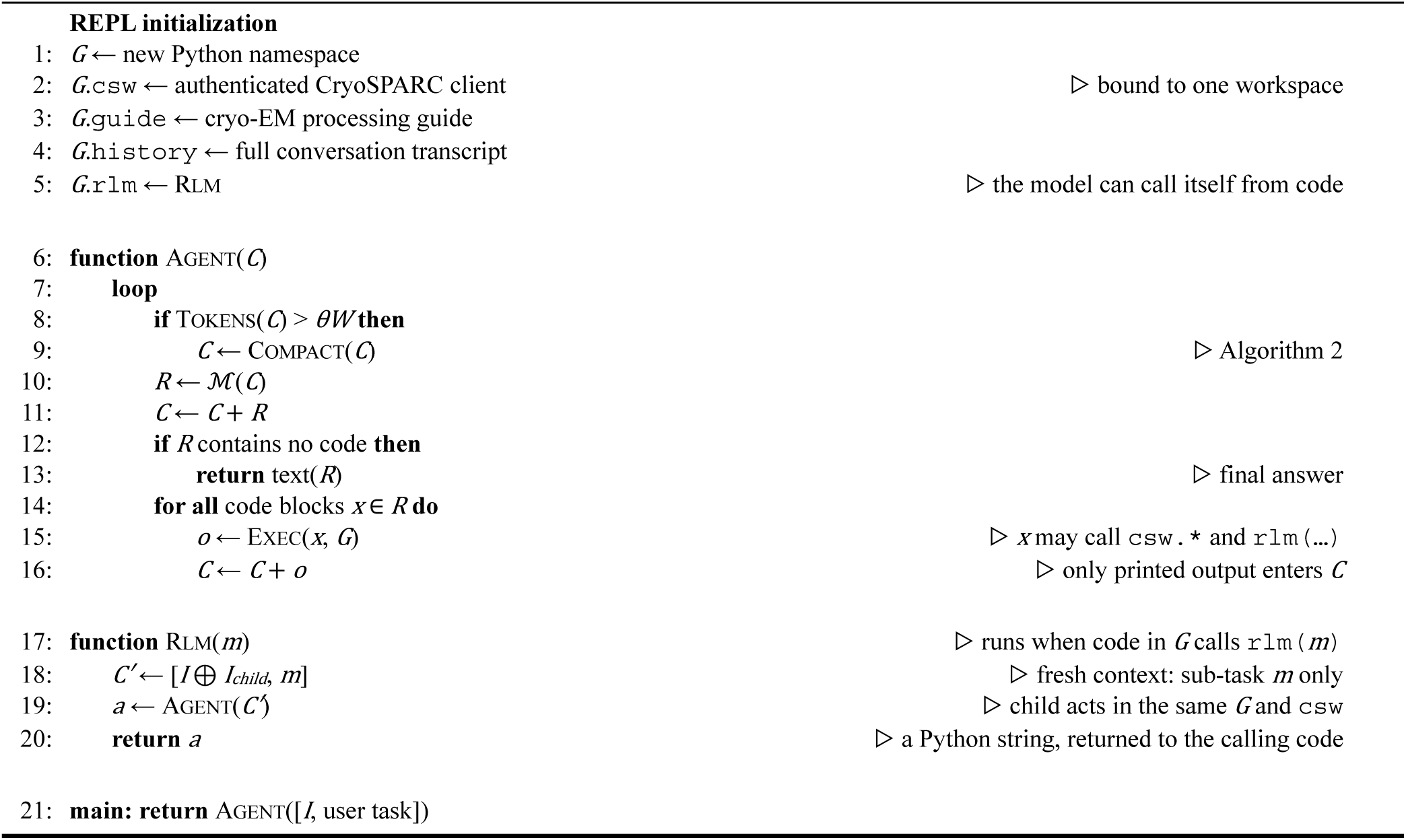

#### Notation

*ℳ* is the language model, *G* is the REPL namespace, and *C* is the active context, the ordered list of messages that the harness sends to *ℳ*. *I* is the system instruction, *W* is the context window in tokens, and *θ* is the compaction threshold as a fraction of *W*. The symbol ⧺ joins message lists and ⊕ joins instructions. A variable that one tool call defines in *G* stays available to all later calls. *R* is a response of *ℳ*, *x* is a code block in *R*, and *o* is its printed output. *m* is a subtask for a child agent, *C*′ is the context of the child agent, and *a* is its answer. In Algorithm 2, *H* is the kept instructions and task, *T* is the most recent turns, and *S* is the summary.

Code can call CryoSPARC through the client *csw*, and it can call rlm(*m*) with a subtask *m*. Rlm starts a fresh context that contains only the instructions, an additional child instruction *I*child, and *m*, and runs Agent on it in the same namespace and CryoSPARC workspace. The child agent returns its answer to the calling code as a Python string, which the parent can parse, print, or pass to other code. Child agents handle iterative tasks that produce long transcripts, mainly rounds of 2D classification and class selection.

They keep these transcripts out of the main context and return only their conclusions. To limit cost, recursion is capped at a depth of 4, 16 recursive calls per session, and 64 model turns per call.

#### Context compaction

A project that runs for several days can exceed the context window of the model. Before each model call, the harness therefore estimates the number of tokens in *C* and, if this exceeds *θW*, compacts *C* with Algorithm 2. *W* is 1,000,000 tokens and *θ* is 0.23.

##### Algorithm 2 Context compaction

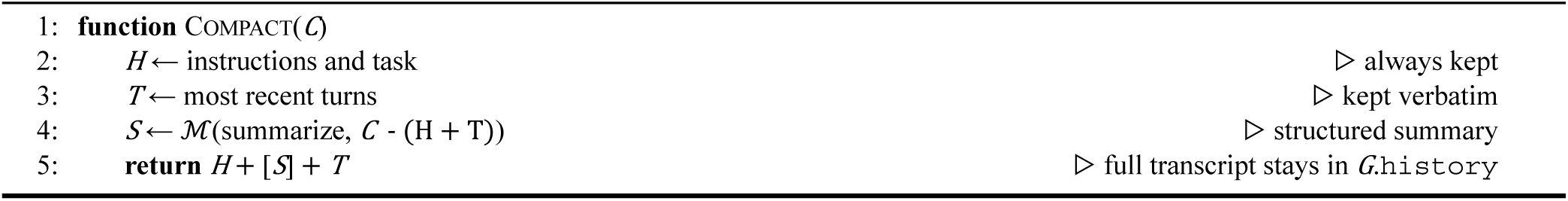

Compaction keeps the instructions, the task, and the most recent turns unchanged and replaces the other turns with a structured JSON summary, which *ℳ* writes from these turns and the previous summary *S*prev. The full transcript stays in *G*.history and in the database, and the agent can search it with memory_search and memory_slice.

#### Sessions

For each run, the researcher gives the data location, the acquisition parameters, a sample description, and the goal, and a launcher creates a dedicated CryoSPARC project and workspace. The database stores the conversation in order, checkpoints of the active context, the compaction summaries, and a record of each recursive call. After an interruption, the harness restores the conversation and the project identity and tells the agent to rebuild its job handles from the saved identifiers and check their status before it submits more jobs. If a tool call was interrupted, the agent is told that the call may have taken effect, so that it does not blindly repeat an action such as a job submission.

### Modeling agent design

#### Agent and loop

The modeling agent is a function-calling agent implemented with the OpenAI Agents SDK ^57^ and GPT-5.5 at the high reasoning setting, with a limit of 500 model turns per run. At each turn, the model reads the conversation, calls one or more tools, receives their outputs, including images when needed, and continues until it produces a final report. Every run receives the same goal: build the most accurate atomic model supported by the map. Inputs are the map, its resolution, one sequence record per chain copy, component stoichiometry, and, when available, an unsharpened map. Runs are independent and cannot reuse models, scores, or CryoZeta jobs from other runs.

#### Tools

The agent has 25 tools for map assessment, handedness determination, model building, validation, visualization, editing, and refinement. render_map_views displays thresholded orthogonal views and reports density connectivity. For handedness, the agent can search and retrieve related PDB structures ^58^, fit them with VESPER, calculate Q-scores ^59^, reflect the map, and compare the two hands. CryoZeta is accessed through tools for job submission, monitoring, and result retrieval, with at most three jobs per run. Model quality is assessed with DAQ-AA, residue-level density support, chain-break checks, and Phenix validation. The agent can also compare DAQ between model versions and inspect map–model overlays. Model editing is performed with edit_model, whereas refinement uses phenix.real_space_refine and Rosetta density-guided FastRelax with agent-selected options and density weights. File-access and shell tools provide additional access to Phenix, Rosetta, and ChimeraX programs.

#### Contour level

The instructions describe three sources of evidence for selecting the contour used by CryoZeta: the isosurface-area curve as contour increases, connectivity of the thresholded density, and enclosed volume relative to the expected assembly volume, estimated at ∼1.21 Å³ per dalton and ∼110 Da per residue. In the reported runs, the agents relied mainly on connectivity and enclosed volume. They selected levels close to the expected molecular volume while allowing for additional density, such as glycans or nanodiscs, and verified that most above-threshold voxels belonged to one connected body.

#### Instructions and outputs

The instructions define the modeling goal and six ground rules. The agent works only within its own workspace, submits CryoZeta jobs only when expected to improve the model, and may use experimental structures as supporting evidence, for example for handedness, but may not copy their coordinates or use structure predictors. Incorrect hand is corrected by reflecting the map rather than the model. The instructions also provide stage-specific guidance for contour and hand selection, candidate evaluation, assembly checks, removal of unsupported residues, refinement, and validation, with typical targets such as MolProbity <2, clashscore <10, and >95% Ramachandran-favored residues. Each run outputs the final model, DAQ and Phenix validation results, a chronological decision log with supporting evidence and alternatives considered, and a structured summary. The agent runs on a CPU node using at most four processors per program, whereas CryoZeta runs on a remote GPU server. A modeling run used 3.3–7.6 million input tokens (92–96% cached) and 19–28 thousand output tokens, about US$3–7 at list prices, and spent 2.0–10.5 h of its wall time waiting for CryoZeta.

## Data and code availability

A binary of the CryoPilot web application will be available at https://cryopilot.vercel.app/.

## Acknowledgements

This work was partly supported by the National Institutes of Health (R35GM158267, R21AI187928) and the National Science Foundation (IIS2211598, DBI2146026, and DBI2422620). PP is supported by a graduate fellowship (T32 GM132024).

## Author contributions

D.K. conceived the study. P.P. designed and developed CryoPilot and the modeling agent, performed all processing runs and analyses, and prepared the figures. P.P and D.K. wrote the manuscript. T.K. advised on 3D reconstruction and instructed P.P. on cryo-EM data processing. D.K. Critically checked and edited the manuscript.

